# AI-driven prediction of SARS-CoV-2 variant binding trends from atomistic simulations

**DOI:** 10.1101/2021.03.07.434295

**Authors:** Sara Capponi, Shangying Wang, Erik J. Navarro, Simone Bianco

**Affiliations:** IBM Almaden Research Center, 650 Harry Rd, San Jose, CA 95120, USA; Center for Cellular Construction, San Francisco, CA, 94158, USA; Graduate Program in Biophysics, University of California, San Francisco, San Francisco, CA, 94158, USA

## Abstract

We present a novel technique to predict binding affinity trends between two molecules from atomistic molecular dynamics simulations. The technique uses a neural network algorithm applied to a series of images encoding the distance between two molecules in time. We demonstrate that our algorithm is capable of separating with high accuracy non-hydrophobic mutations with low binding affinity from those with high binding affinity. Moreover, we show high accuracy in prediction using a small subset of the simulation, therefore requiring a much shorter simulation time. We apply our algorithm to the binding between several variants of the SARS-CoV-2 spike protein and the human receptor ACE2.

## 1 Introduction

The ongoing COVID-19 pandemic, caused by the infection with the RNA betacoronavirus SARS-CoV-2, has already resulted in over 180 million cases and 3.98 million deaths around the world [1]. As the global response to the pandemic moves from non-pharmaceutical (e.g., social distancing, periodic lockdowns, mandate of personal protective equipment) to pharmaceutical (e.g., vaccination) forms of intervention, important questions arise about the evolution of the virus and the emergence of genetic variants which may escape the drug. Coronavirus viral particles contain four structural proteins [2]: the glycoprotein spike (S), the membrane (M), the envelope (E), and nucleocapsid (N) proteins. The S protein (Fig. 1A), which is the main focus of much of the ongoing research, is a trimeric class I membrane fusion protein responsible for binding the host cell receptor and triggering the fusion between the viral membrane and the host cell membrane [3–6]. The S protein exists in a prefusion metastable state and undergoes large conformational changes transitioning to a stable postfusion conformation once it binds to a host cell receptor [3, 6–8]. Each S protomer (see Fig. 1A) comprises two functional subunits: S1 contains the receptor binding domain (RBD) and is responsible for binding the host cell receptor; S2 contains the fusion machinery and is responsible for the membrane fusion between the virus and the host cell. The first cryo-EM structures of the SARS-CoV-2 S protein in the prefusion conformation, published in March 2020, revealed fundamental structural details providing the basis for identifying conserved and accessible epitopes for future antibody isolation and vaccine design efforts [9, 10]. In addition, several structures of the SARS-CoV-2 RBD in association with ACE2, published between March and May 2020, helped uncover the molecular mechanisms of the interaction between the RBD and ACE2 and to identify residues that are crucial for the binding process [11–14].

**Fig. 1.**
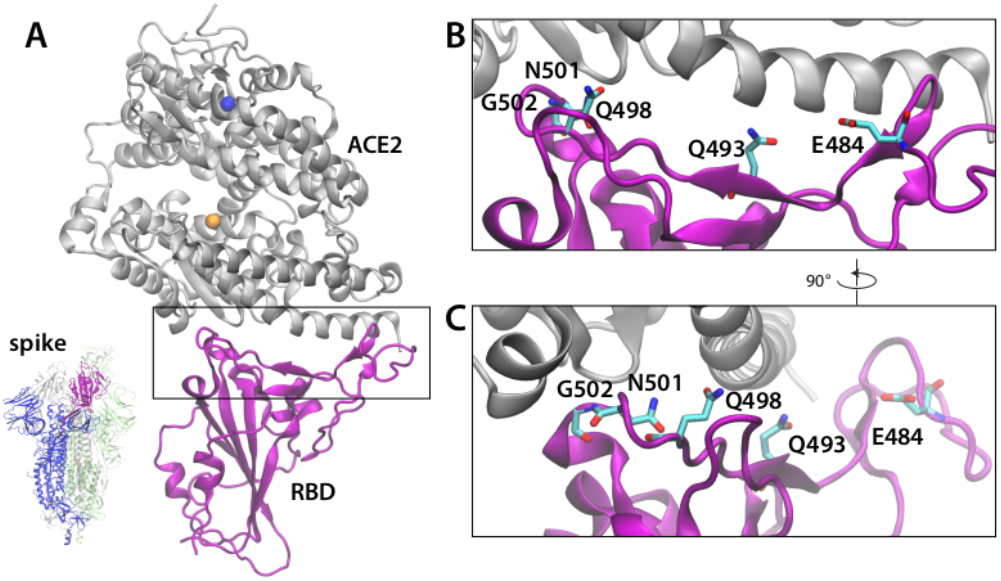
**A**. Representative snapshopt of the chimeric spike receptor binding domain (RBD) (purple) in association with ACE2 (gray). Proteins are represented in New Cartoon format. Zinc (orange) and chloride (blue) ions are represented in van der Waals format. Bottom left: representation of the spike trimeric complex from [9], one chain is blue with the RBD colored purple, the other 2 chains are transparent gray and green for clarity. **B-C**. Closeup view of the contact region between RBD and ACE2. The non hydrophobic, non aromatic residues that are mutated and studied in this paper are indicated. They are represented in licorice format colored according to the ion type (red for oxygen, cyan for carbon, blue for nitrogen). Hydrogen atoms are not displayed for clarity. **C**. Same representation of panel **B** rotated of 90°.

Being the RBD the most variable part of the viral genome [15], several studies have focused on understanding the effects mutations have on the binding affinity to ACE2 [16–18]. Atomistic molecular dynamics (MD) simulations have been used since early in the pandemic to study and visualize the binding between the S and the ACE2 receptor and explore potential targets for drugs [19–30]. Three contact regions (CR1, CR2, and CR3) have been identified as the key binding points between RBD and ACE2 [23]. CR1 and CR3 are located at the ends of the RBD-ACE2 interface while CR2 refers to the interface middle region. In CR1 and CR2, the hydrophobic contacts between RBD and ACE2 mediate the interaction, whereas CR3 is characterized mostly by hydrogen-bond (H-bond) forming residue interactions. An important limit of atomistic MD simulations is the often times high computational burden necessary to carry out the simulations to an extent that is meaningful to answer biological questions. To estimate the free energy of binding between the RBD and ACE2, and more in general between two proteins, one would need to carry out one simulation of the association process starting from separate protein molecules, and another simulation sampling the dissociation process. However, the association and dissociation rates can be very slow, much beyond the current atomistic MD simulation accessible time scale, making it hard to estimate directly the binding free energy from sampling association and dissociation events *in silico*. Therefore, several theoretical approaches have been developed to solve this issue. Among others, Markov state modeling [31–33], Hamiltonian replica exchange methods [34], weighted ensemble approaches [35, 36], or adding a bias potential (umbrella sampling) to guide the association or dissociation process along a progress coordinate [37, 38] are among the most common tools to study the binding process between two molecules. Often, free-energy perturbation (FEP) calculations are used to investigate the energetic contribution of the mutated amino acid to the proteinprotein binding interaction [22, 24, 39]. Once an approach has been selected, the following biological question arises: how can we probe efficiently *in silico* the effects of site mutations on the protein-protein interaction and on the binding affinities? The answer to this and other important questions is increasingly relying on machine learning (ML) methods, whose application in computational biochemistry is gaining momentum.

Indeed, artificial intelligence (AI) has recently emerged as a potential accelerator of simulations of dynamical systems, including MD simulations [40–44]. For MD simulations, the majority of ML methods have focused on learning the force field [41], that is, the complex set of rules and parameters governing the interaction among atoms and thus between molecules. Recently, an increasing amount of works devoted to the prediction of experimentally measurable quantities from MD has emerged [40, 45–47]. Specific to SARS-CoV-2, in a recent paper, MD simulations and ML methods are combined with FEP calculations to investigate the impact of individual residues on binding to ACE2 in SARS-CoV-2 compared to SARS-CoV [22].

In our work we present a neural network-based method to predict molecular binding trends, that is, whether or not the binding affinity of a target molecular interaction is higher or lower than a reference interaction, and reduce the computational burden of the molecular simulations. Rather than learning the sequence to structure relationship as most AI methods in this space do [48], our method learns the relative atomic distance between molecules over the simulation time and uses this information to predict the molecular association or dissociation trend. We apply our method to evaluate binding between mutants of the SARS-CoV-2 RBD and the ACE2 receptor (see Fig. 1). We demonstrate that our method is reliable and accurate for predicting binding affinity trend for non hydrophobic mutations occurring at the RBD-ACE2 interface, and has limited applicability to non hydrophobic mutations. Our goals are twofold: First, we aim to demonstrate that the combination of AI and MD simulations allows for the prediction of the binding affinity trend between two molecules. While it is difficult to measure affinity through MD for systems of the size of the RBD-ACE2 complex, we prove that, if that information is experimentally known for a set of mutants, a neural algorithm is able to predict the correct affinity trend of an unknown mutation. Secondly, we prove that our AI algorithm can be used to obtain reliable information about binding affinity trends from short MD simulations, thereby significantly reducing the time to obtain the prediction.

Our paper is structured as follows: The next section introduces our MD and AI methodologies, as well as the data. Next we report the results of our analysis. Finally we discuss the potential impact of our work and future directions.

## 2 Data and Methods

### 2.1 Reference data

We obtain binding affinity for all mutations of the S protein RBD–ACE2 receptor from the recent literature [16]. The non hydrophobic mutated residues are represented in Fig. 1B-C and a list is reported in Table 1 according to their decreasing value of binding affinity: six mutations exhibits higher affinity and six lower affinity than the wild type (WT), which we use as reference. In Table 1 Δ log_10_(*K_D,app_*) quantifies the relative affinity value compared to the unmutated SARS-CoV-2 RBD, with positive values indicating stronger binding, and negative values indicating weaker binding. We choose two medically important mutations, N501Y (present in the Alpha variant, formerly known as the English variant) and E484K (present in the Gamma variant, formerly known as Brazilian variant), two mutations which are present in SARS-CoV, N501T and Q493N, and two mutations from the bat coronavirus RaTG13. Additionally, we randomly selected five additional variants according to their affinity, for the purpose of balancing our training set. Additionally, we selected four hydrophobic, non aromatic sites (A475, L455, L492, V503) and mutated those residues to hydrophobic, non aromatic residues, for a total of nine additional simulations. According to Ref. [16], the RBD non aromatic hydrophobic amino acids at the RBD–ACE2 interface are only two: V503 and L455. The complete list of the hydrophobic mutations is reported in Table A1, along with the origin of the mutation, the binding affinity value, and the simulation length.

**Table 1.** List of the mutations of the spike RBD studied in this paper and used to train the CNN. The second column reports the origin of the mutation and the third column the binding affinity extracted from Ref. [16]. The mutations are listed according to decreasing values of the binding affinity.

| Mutation | Origin | $\Delta \log_{10}(K_{D,app})$ |
| --- | --- | --- |
| N501Y | Alpha variant | 0.24 |
| Q498Y | SARS-CoV | 0.16 |
| N501V | - | 0.15 |
| Q493Y | Bat RaTG13 | 0.12 |
| N501T | SARS-CoV | 0.1 |
| E484K | Gamma variant | 0.06 |
| N501S | - | -0.13 |
| Q493N | SARS-CoV | -0.21 |
| Q498N | - | -0.5 |
| Q498K | - | -2.26 |
| N501D | Bat RaTG13 | -2.42 |
| G502P | - | -4.55 |

### 2.2 Molecular dynamics simulations

To carry out the simulations of the SARS-CoV-2 RBD in association with the ACE2 receptor, we used the atomic coordinates extracted from the X-ray crystal structure of the SARS-CoV-2 chimeric RBD in complex with the human receptor ACE2 (PDB ID 6VW1) [49] (Fig. 1A). The structure has a resolution of 2.68 Å and is a chimera of the RBD from SARS–CoV and the receptor binding motif (RBM) from SARS–CoV–2, with the exception of an RBM loop. For our work, we considered this structure as reference for the WT binding affinity, in analogy with the WT used as reference for the experimental work [16]. We used Solution Builder from CHARMM-GUI [50, 51] to model the disulfide bonds, to solvate the system, add the zinc and chloride ions present in the crystal structure, and add the potassium ions to maintain the system charge neutrality. The final system contains approximately 132000 atoms, with ≈ 40000 water molecules.

From the equilibrated structure of the RBD-ACE2 system, we extracted the last configuration and used the Mutator Plugin from VMD software [52] to build the mutated systems used for this study and reported in Table 1 and Table A1, with the charged and polar mutations represented in Fig. 1B-C. We visually inspected the mutated protein and used VMD software [52] to model the disulfide bonds, to solvate the system, add the zinc and chloride ions present in the crystal structure, and add the potassium ions to maintain the system charge neutrality. The procedure to minimize and equilibrate the system was the same for WT and all mutations, and it is described in Section 2.2.1. The simulations had variable lengths, ranging from 160 to 280 *ns* (Table A1, A2).

All simulations were performed using NAMD 2.14 [53, 54] with the CHARMM36m force field for the protein and ions [55] and the TIP3P model for water [56]. We used a Langevin dynamics scheme to keep the temperature constant at 303.15 K and an anisotropic coupling in conjunction with Nosé-Hoover-Langevin piston algorithm to keep the pressure constant at 1 atm [57, 58]. Periodic boundary conditions were applied in three dimensions. We employ the smooth particle-mesh Ewald summation method to calculate the electrostatic interactions [59, 60] and the shortrange real-space interactions were cutoff at 10 Å using a switching function between 8 Å and 10 Å. The equations of motion were integrated with a time step of 4 *fs* for the long-range electrostatic forces, 2 fs for the short-range nonbonded forces, and 2 *fs* for the bonded forces by means of a reversible, multiple time-step algorithm [61]. The SHAKE algorithm [62] was used to constrain the length of the bonds involving hydrogen atoms. The simulations were visualized using VMD software [52].

#### 2.2.1 System Equilibration Protocol

We used the following procedure to minimize and equilibrate all the systems. To minimize the system, we used the conjugate gradient algorithm for 8,000 steps and then gradually heated the simulated cell from 25 K to 300 K. To equilibrate the positions of ions, water molecules, and protein complex, harmonic restraints were used in a series of six consecutive simulation runs of 1 *ns*, as follows: we ran 1 NVT (constant number of particles N, volume V, and temperature T) and 5 NPT (constant number of particles N, pressure P, and temperature T) equilibration runs. In the first equilibration run, restraint force constants of 20 kcal mol^−1^ Å were applied to the protein backbone atoms and of 5 kcal mol^−1^Å^−2^ to ions and water. In the subsequent runs, the restraints on the protein backbone were decreased to 10, 8, 4, 2, and then to 1 kcal mol^−1^Å^−2^, while restraints on ions and water were decreased to 2, 1, 0.5, 0.1 and then 0.01 kcal mol^−1^ Å^−2^, respectively. After the last 1 *ns* equilibration run, we began the production runs. The simulations were carried out under NPT conditions and the coordinates were saved every 20 *ps*.

#### 2.2.2 Dynamics of the systems from MD simulations

For all systems, from the equilibrated structure we carried out simulations in the NPT ensemble (Table A1, A2), monitored visually the simulations, and calculated the root mean square deviations (RMSDs) of all the C*α* carbon atoms belonging to both proteins and of the C*α* carbon atoms belonging to the RBD and ACE2 contact area (see Figures A1, A2, and A3; Section Appendix A). The contact area appears to be rather rigid in almost all systems with C*α* RMSDs < 2Å. In the mutation N501D the RMSD value reaches ≈ 3Å between 180 *ns* at the end of the simulation, also in the L492 system the RMSDs increase to ≈ 3Å at around 100 *ns* but after that they decrease to ≈ 2Å. In V503A we observe a detachment in CR1 and CR2 explaining the increase of the RMSD up to ≈ 4Å at the end of the simulation. In few simulations and at different time points of the simulation, we observe escape of the Cl ion from the ACE2 binding pocket. We do not observe reduced stability due to loss of Cl, and verify that it does not impact the RMSD throughout the affected simulations.

#### 2.2.3 Training set preparation

The result of the MD simulations is a time series of molecular snapshots. We use the following procedure to convert the data into a training set for our machine learning algorithm (Fig. 2). From the long simulation of the RBD in association with ACE2, we extracted the residue identification number of the C*α* carbon atoms of the RBD that happen to be within 6 Å of ACE2 receptor at least for a single frame during the whole simulation length. We repeated the same analysis for ACE2, obtaining two lists of residues (55 amino acids for the RBD and 62 for ACE2 respectively) that represent the amino acids found in one protein within 6 Å from the other and viceversa (Fig. 2A); for the complete list of amino acids see Appendix A. We called this region as RBD and ACE2 *contact area*. For each system, we calculated the distances among all the C*α* carbon atoms of the residues belonging to the two lists for each frame. For this, we used the contact and distances modules from MDAnalysis [63, 64]. Finally, we used the matrices of the distances to construct images that can be interpreted by our machine learning algorithm as follows (Fig. 2B).

**Fig. 2.**
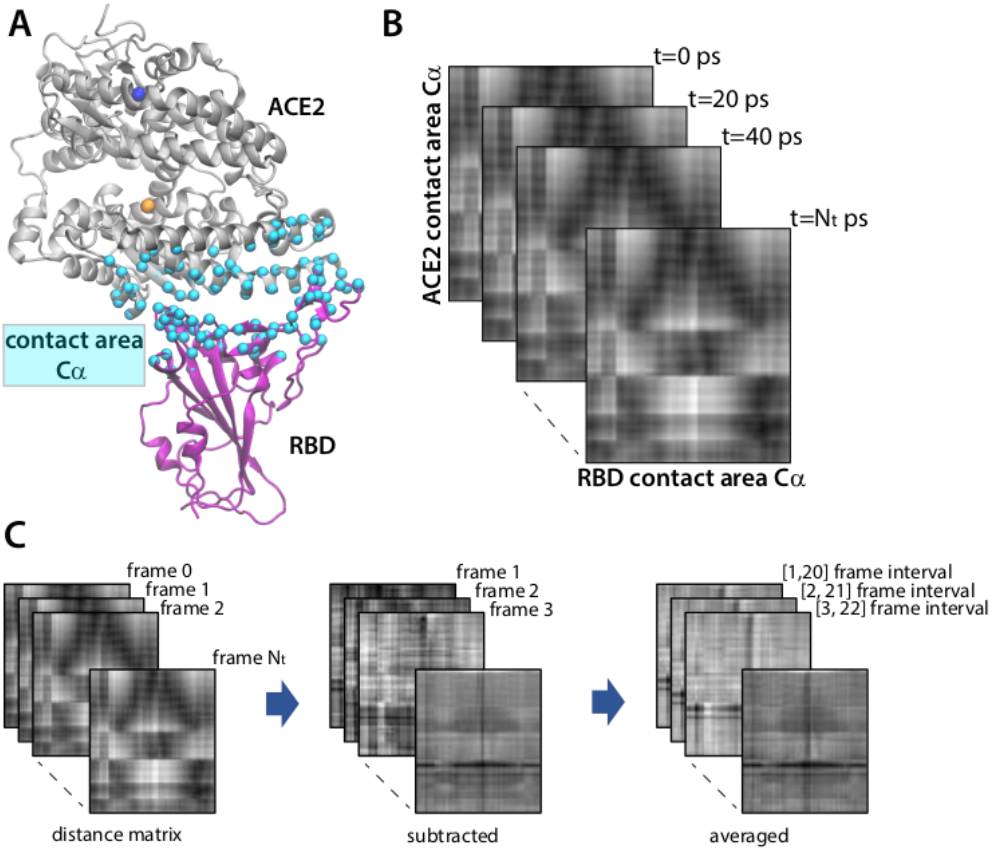
Conversion of the MD data to input matrices for a neural algorithm. **A**. Representative snapshot of the chimeric RBD-ACE2 complex indicating the C*α* carbon atoms used to calculate the distance between the RBD and ACE2. The color code is as in Fig. 1 and the C*α* carbon atoms are represented in cyan, van der Waals format. The distances between the atoms are then represented as matrix, are normalized by each of their minima and maxima, and converted to gray-scaled images. **B**. Representative distance matrices of the chimeric RBD-ACE2 complex simulations taken at different steps of the simulation showing the effect of data preprocessing. **C**. Representation of the data processing to obtain machine learning inputs. We first subtract the distance matrix at *t* = 0 from the distance matrices from the simulations at each time steps (central panel). We then use a moving average to average the subtracted matrices and use these as input to the algorithm (right-hand panel).

We recorded the distance matrices every 20 *ps* to up to 160*ns*, i.e., 8000 molecular snapshots were collected and preprocessed. Simulations of under 160*ns* were not sufficiently long to directly reveal the binding trend. The dissociation constants of typical protein–protein complexes are in the nanomolar range and the dissociation rate can be of the order of minutes or longer, much beyond the timescale of current MD simulations. The time evolution of the distance matrices shows very similar patterns for all mutations (Fig. 2B and also the supp videos). Therefore, we perform the following procedure to highlight the difference in the molecular dynamic due to the mutations (Fig. 2C). The signals are buried in the structural restrictions of the positions of the amino acids. In order to observe how the atoms move due to the contact between RBD and ACE2, we first subtract from all matrices the initial frame of coordinates after completion of minimization and equilibration procedures, which we indicate as *t* = 0 (see 2.2.1). We then average the signal using a moving average of 20 frames (or 400 *ps*) to reduce noise (Fig. 2C). We then normalize the averaged matrices using min-max normalization to make sure all values in the matrices are between 0 and 1. Finally, we converted them to gray-scale images and use them as neural network inputs for training, validation and testing. In Fig. 3 we show a representative image for each of the polar and charged mutations analyzed. The training and validation sets were built from the polar and charged mutations using a randomly chosen 90/10 split of all input images from three mutations showing higher affinity than the reference (namely, N501Y, N501V, and N501T) and three mutations showing lower affinity than the reference (namely, N501S, Q498N, and N501D). All images from the remaining six mutations were used for testing. Such trained CNN was then used for testing also on hydrophobic, non-aromatic mutations.

**Fig. 3.**
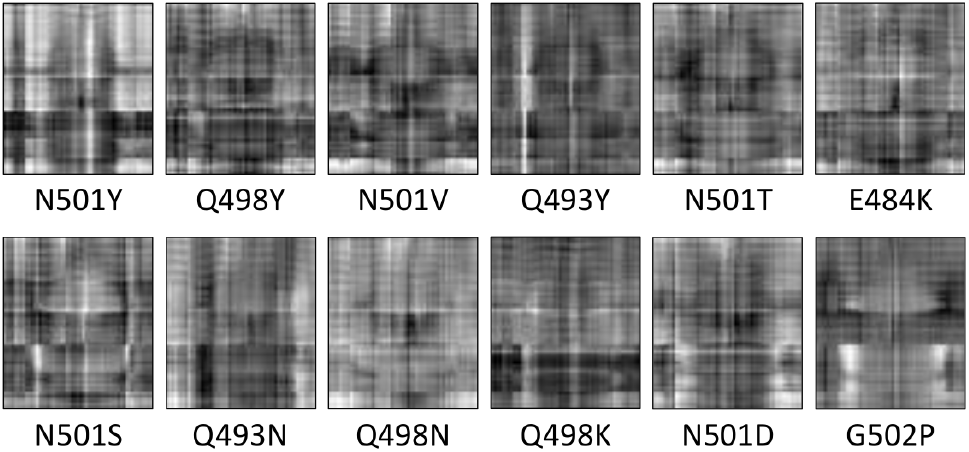
Sample gray-scaled images representing the averaged matrices for some of the mutations studied in this paper at *t* = 20 *ns*.

### 2.3 Neural Network Framework

#### 2.3.1 Neural network Architecture

In this work we used a Convolutional Neural Network (CNN) implemented in Python 3.7.8 and TensorFlow v2.4.1. The neural network architecture is illustrated in Fig. 4A. The neural network uses 62 × 55 grayscale images as inputs, and outputs two values corresponding to the probabilities of higher and lower affinity compared to the WT (*P*_1_ and *P*_2_, respectively). Between the input and output layers, we have 2 convolutional layers and 2 maeonximum pooling layers. Convolutional layers detect spatial correlations in input data and maximum pooling layers down-sample input data to reduce dimensionality and the number of model parameters. We used RandomNormal initializer and ReLU activation function for the convolutional layers. The outputs of the second maximum pooling layer was then flattened. We used dropout regularization to the flattened array with dropout rate equal to 0.5 to prevent over-fitting. The flattened layer then connected to a dense layer containing 5 nodes. We used softmax activation function to connect this dense layer and the final output layer to convert the outputs into categorical probabilities, i.e. *P*_1_ + *P*_2_ = 1. If *P*_1_ > *P*_2_, the input will be categorized as having higher or lower binding affinity than WT (Fig. 4C). For training, we used sparse categorical cross entropy loss and Adam optimization algorithm with the initial learning rate equals 0.0001, and training over 30 epochs.

**Fig. 4.**
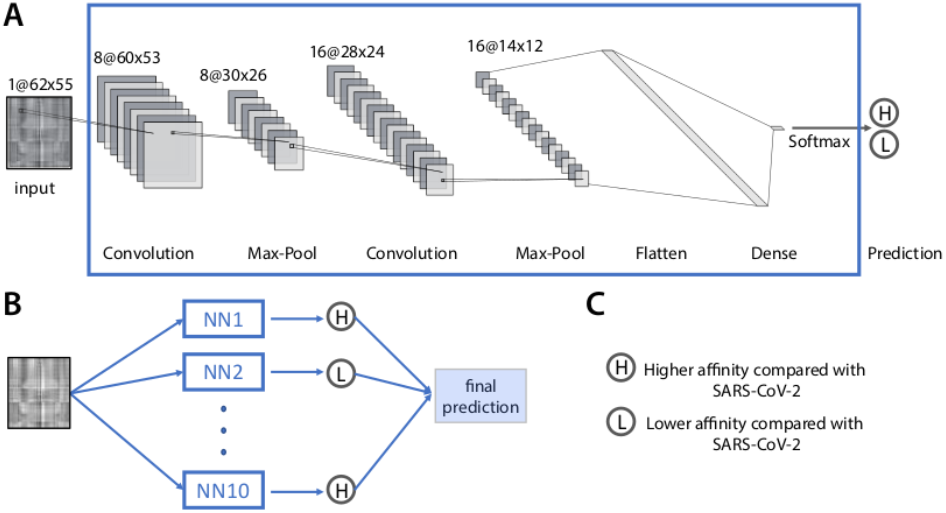
Structure of the convolutional neural network and ensemble prediction used in this study. **A**. The convolutional neural network architecture. **B**. The ensemble prediction consisting of *N* independently trained convolutional neural networks. The final prediction is based on the simple majority of all neural networks. **C**. Definition of the output categories.

#### 2.3.2 Ensemble Predictions

Due to the stochastic nature of network initialization and dropout, as well as the availability of a limited training set, every neural network is unique in terms of the parameterization of the network connections [43, 65]. To mitigate the potential impact of this issue on the classification, we implemented an ensemble decision method to get consensus prediction: for each mutation, we train ten identical neural networks. We then use a simple majority to assign the final class to the mutation (Figure 4B). This method has been proven to further reduce prediction errors and bias and increase accuracy and reliability, especially with limited training data [43, 65, 66].

## 3 Results

### 3.1 An ensemble of CNNs can reliably predict binding trends of polar and charged amino acids

The results of our ensemble procedure are reported in Tables B1 to B6, and summarized in Table B7. Every row represents a different network run, while every column is the result of testing using only a subset of the data. The row labeled FP is the final prediction for the ensemble, summarized by the accuracy of prediction in the row labeled PPC. We first focus on the predictions on non hydrophobic residues, using the whole MD simulation for testing (last column in the Tables, *t* = 160*ns*).

Nine out of ten trained neural networks can predict the six mutations in the test set with 100% accuracy, while one neural network can predict four out of six mutations in the test set, with an overall average accuracy of 0.9667. Thus, we conclude that our methods is optimal in predicting binding trend of non hydrophobic mutations with respect to a reference with high accuracy. Moreover, by using the ensemble prediction method, we show that we successfully eliminate the possibility of an incorrect prediction due to faulty training, and increase our confidence when making a call on a novel mutation.

We then test our method on hydrophobic, non-aromatic residues (table A1). Our method can predicts with high accuracy the binding affinity of V503I, V503A, and L455I, with less precision that of L492I, and fails in predicting L455A, L455V and 3 mutations of the A475 residue (see Table B8). We rationalized our result by considering that the CNN training set only includes mutations of charged and polar amino acids localized in the contact region CR3 [23], which is at one end of the RBD-ACE2 interface. The trained CNN is able to predict the binding affinity trend of non hydrophobic mutations located in CR1, CR2, and CR3 and of a hydrophobic mutation in CR3 with high accuracy, but is less precise or fails in predicting hydrophobic mutations localized in CR2 and CR1. Including one hydrophobic mutation of CR1 in the training set (e.g. A475P) improves the predictive accuracy for other CR1 hydrophobic mutations (e.g. A475L) but decreases the accuracy of the H-bond forming mutations in CR3 (e.g. Q498Y) and that of a hydrophobic mutation in CR2 (e.g. L455V). While more work is necessary to address this issue, the inclusion in the training set of examples of mutations from all contact regions is potentially necessary for the generalization of the methods to hydrophobic mutations for this particular system.

### 3.2 An ensemble of CNNs can accurately predict binding trends of polar and charged amino acids using shorter MD simulation times

We now discuss the problem of how much MD data is needed to correctly predict whether a non hydrophobic mutation has higher or lower than WT affinity using a CNN trained in the way we described. As stated above, MD simulations of minutes or longer are needed to observe binding or dissociation. In the previous sections, we showed that, when the ground truth is known, simulations of 160 *ns* of six systems of the RBD-ACE2 complex size are sufficient to train neural networks which can reliably predict binding trends for unknown mutations. However, performing MD simulations of 160*ns* of 12 or more systems can be still a challenging task in the absence of adequate computing capabilities. This can limit the ability to simulate important molecular processes in situations where rapid responses would be necessary. To give an estimate of the necessary resources, on the Artificial Intelligence Multiprocessing Optimized System (AiMOS) supercomputer using NAMD 2.14 [53, 54] (see Method) we are able to simulate ≈ 33*ns* per simulation per day using 16 2.5 GHz Intel Xeon Gold CPUs, 768*Gb* ram, and 8 NVIDIA Tesla V100 GPUs (from the benchmark test simulations we carried out it resulted that 8 GPUs was the best efficient usage of AiMOS resources for our systems), while we are able to simulate ≈ 7*ns* per day on a cloud bare metal server consisting of 2.4*GHz* Intel Xeon CPUs, 128*Gb* ram, and 4 Nvidia K80 GPUs. In this section, we test whether or not it is possible to use our CNN to accurately predict binding trends using shorter MD simulations. We varied the duration of the data extracted from the simulations from 20 *ns* to 160 *ns*, with increments of 20 *ns*. Our results are reported in Tables B1 to B6 for single mutation predictions, and summarized in Table B7 and Fig. 5.

**Fig. 5.**
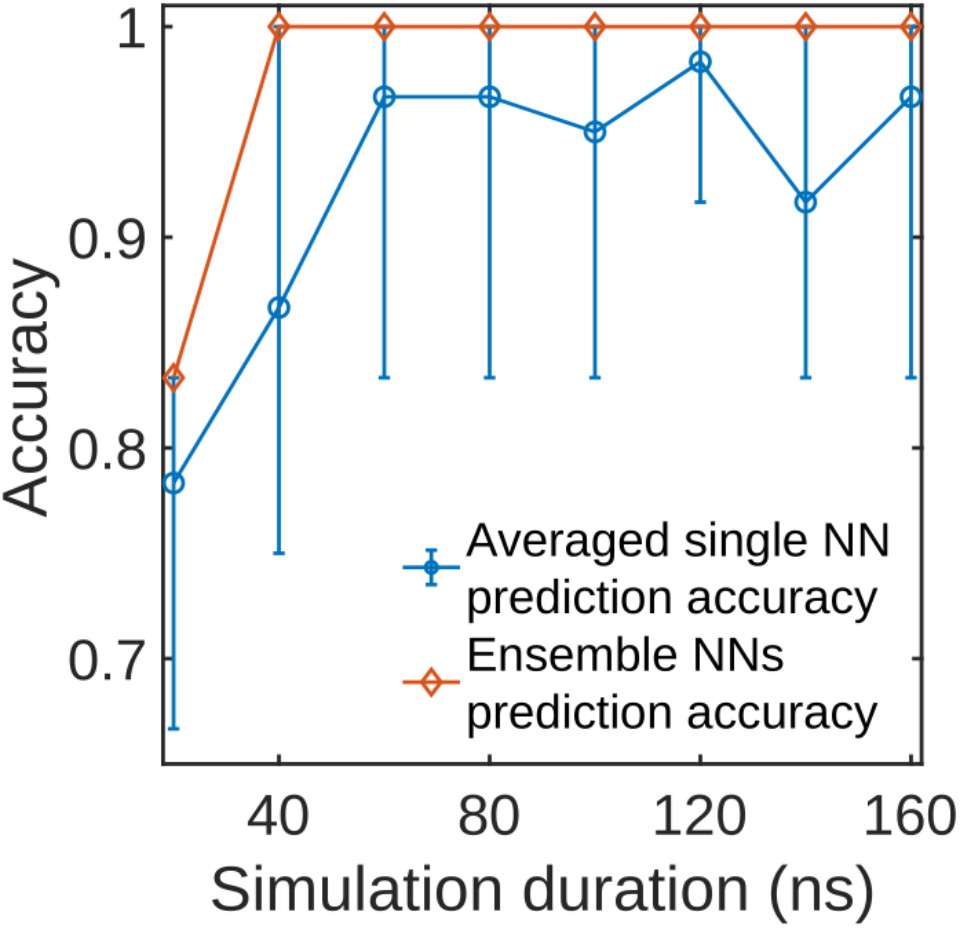
The overall test accuracy of CNNs varies with the simulation time. The blue dots are the average prediction accuracy from ten independently trained CNNs. The error bars represent the 10^*th*^ percentile and 80^*th*^ percentile. The orange diamonds represent the prediction accuracy of the ensemble prediction, voted by 10 NN assembles.

Figure 5 reports the average prediction accuracy of ten independently trained neural networks as well as the ensemble accuracy as a function of simulation time. The error bars for the averaged predictions represent the 10^*th*^ percentile and 80^*th*^ percentile of the prediction accuracy (the 90^*th*^ percentile is equal to the 80^*th*^ percentile, see Table B7 for details). We can see that the average prediction accuracy for independently trained CNNs are consistently over 90% with the simulation duration equal or longer than 60 *ns*. Based on the prediction accuracy of this ten independently trained neural networks, the optimal simulation time for the averaged single neural network predictions for this specific system is 120*ns* (see Table B7 for details). Taking advantage of the ensemble prediction method, the accuracy can be as high as 100% with simulation duration as short as 40*ns*. The ensemble prediction significantly increases the prediction accuracy and reliability while allowing for a 4*x* shorter simulation time. Furthermore, we observe that some mutations such as E484K, Q493N, and G502P can be correctly classified with high confidence (> 75%) with only 20 *ns* of simulation. This result can be explained by the fact that, during protein-protein interactions, structural modifications following mutations affecting H bond networks can be observed at relatively short times (see for instance [67–69] and references therein). The disruption of an H–bond and/or a salt bridge affects greatly the amino acid side chains, but the distance matrices based on which the images to train and test our CNN are generated are calculated between the C*α* carbon atoms of the amino acids of the RBD and ACE2 belonging to the contact area (as defined it in the Method section). Therefore, our CNN is more sensitive to perturbations occurring at the level of the backbone of the protein rather than to those occurring at the side chain level. In other words, for some mutations, it takes longer to propagate the perturbation of breaking an H bond from the side chain to the protein backbone, which results in the observed accuracy at short simulation times.

Moreover, we suspect that the molecular snapshots of E484K, Q493N, and G502P have high similarity to the snapshots included in the training set, while mutations such as Q498K and Q498Y that are relatively more challenging for the network to classify at shorter time horizons may show greater differences with respect to those included in the training set. A further investigation on the features learned by the CNN would help understanding the different performance exhibited by the CNN in classifying distinct mutations. Also, a training set composed of a more diverse set of mutations will possibly result in more consistent predictions with respect to data extracted form short time window simulations. Nonetheless, our results prove that our CNN is capable of predicting binding affinity trends with high accuracy at very short simulation times when the training set is composed of only a handful of non hydrophobic mutations.

## 4 Conclusions and future directions

The emergence of viral variants in a pandemic epitomizes the evolutionary arm race between the virus and the host immune system. As vaccines are deployed to a growing fraction of the population, questions regarding drug resistance and escape will likely dominate the public and scientific discourse of the near future. Therefore, time is of the essence to determine whether a specific genetic variant has a higher or lower potential to bind to the human cellular receptor. MD simulations are an important tool to address these issues. Methods that accelerate discovery are now necessary for these computational approaches to yield timely responses. Once trained, our CNN takes less than a second to perform a classification task.

While our methods excels on the inference of the binding affinity trends for charged and/or polar amino acids, it only partially succeeds when applied to hydrophobic mutations. An explanation for this limitation may be related to the position, the characteristic of the mutations used for the training set, and the sparseness of the input space. Our training data set includes 6 non hydrophobic mutations localized in CR3, whereas for the second training data set we used a non aromatic, hydrophobic amino acid located in CR1 in addition to the previous ones. A more generalizable predictor needs a training data set constituted of mutations of different type of amino acids such as aromatic, hydrophobic, charged, and polar, and localized over the entire interface between the two proteins (for RBD-ACE2 interface these mutations used for the training set would be in the three contact regions CR1, CR2, and CR3). We plan to improve and further validate our method by adding to the training set more mutations of different type of amino acids localized in different regions. In addition, we also plan to combine the images, i.e., the distance matrices, with other progress coordinates that characterize further the binding process between the two proteins. This will help the learning process of the CNN and will result in better and consistent accuracy in the predictions.

Beyond the application presented in this paper, we believe that a method like the one we propose may be applied generally to other problems where assessing binding affinity trends using only MD simulations may be important, like drug design or cellular engineering. The generalization of these methods is an important argument of research which we are planning to explore in the future. Concurrently, we realize that our methods does not provide an exact measure of the affinity. Recent papers have proposed methods capable of achieving that goal, with mixed but encouraging results [70]. Another future objective of our research is to generalize our method to obtain estimates of binding affinity using limited MD simulations.

## Appendix A: MD simulation details

In table A1 we report the list of the hydrophobic, non-aromatic mutations we carried out along with the mutation origin, the binding affinity extracted from Ref [16], and the length of the simulations.

**Table A1.** List of the mutations of the spike RBD hydrophobic amino acids studied in this paper. The second column reports the origin of the mutation, the third column the binding affinity extracted from Ref [16], and the fourth column the simulation length. The mutations are listed according to decreasing values of the binding affinity.

| Mutation | Origin | $\Delta \log_{10}(K_{D,app})$ | length (ns) |
| --- | --- | --- | --- |
| V503I | SARS-CoV | 0.05 | 187.8 |
| L492I | - | 0.03 | 261.24 |
| L455I | - | -0.01 | 272.24 |
| V503A | - | -0.06 | 188.18 |
| A475V | SARS-CoV | -0.14 | 226.6 |
| L455A | - | -0.43 | 280.72 |
| L455V | - | -0.73 | 280.98 |
| A475L | - | -1.27 | 279.28 |
| A475P | SARS-CoV | -1.62 | 281.02 |

In table A2 we report the length of the simulations we carried out for the polar and charged mutated residues. To monitor the simulations, we calculated the RMSD of the C_*α*_ atoms of the whole proteins and of the contact area between RBD and ACE2 (Fig. A1, A2, A3). For the contact area of the RBD, we considered the C_*α*_ of the following residues: 403, 405, 406, 408, 416 to 418, 421, 437, 439, 440, 444 to 449, 452, 453, 455 to 458, 472 to 478, 484 to 508. For the contact area of ACE2, we considered the C_*α*_ of the following residues: 19 to 39, 41, 42, 45, 48, 49, 61, 72, 75 to 84, 97, 323 to 332, 351 to 357, 386 to 390, 393.

**Table A2.** List of the simulations carried out in this work and of the corresponding length.

| simulation | length (ns) |
| --- | --- |
| RBD-ACE2 | 270 |
| N501Y-ACE2 | 187 |
| Q498Y-ACE2 | 206.84 |
| N501V-ACE2 | 204.86 |
| Q493Y-ACE2 | 197.28 |
| N501T-ACE2 | 186.58 |
| E484K-ACE2 | 204.56 |
| N501S-ACE2 | 205.2 |
| Q493N-ACE2 | 197.18 |
| Q498N-ACE2 | 198.84 |
| Q498K-ACE2 | 204.46 |
| N501D-ACE2 | 210.44 |
| G502P-ACE2 | 160.58 |

## Appendix B: Details of CNN predictions

Table B1, B2, B3, B4, B5 and B6 list the predictions from 10 CNNs for mutation Q498Y (H), Q493Y (H), E484K (H), Q493N (L), Q498K (L) and G502P (L) respectively. These 10 neural networks have the same architecture and are independently trained. The neural network weights are randomly initialized from a normal distribution. Neural network prediction is the probability of higher affinity (*P*_1_) and the probability of lower affinity (*P*_2_) compared to the WT, with the constraint of *P*_1_ + *P*_2_ = 1. H and L is the neural network prediction. H represents higher the WT affinity (*P*_1_ > *P*_2_) and L represents lower than WT affinity (*P*_1_ < *P*_2_). The value in the brackets is the probability value of the predicted affinity. (*P*_1_ for H and *P*_2_ for L). FP is short for final prediction and PPC represents the percentage of CNNs predicting correctly. When training for much smaller data size (simulation ends at 20*ns* and 40*ns*), we trained the CNNs over 60 epochs, while for the rest 30 epochs are used.

Table B7 lists the prediction accuracy of 80 CNNs for all 6 test mutations with different simulation duration, i.e., Q498Y, Q493Y, E484K, Q498K, Q493N and G502P. This table also contains the average accuracy of 10 CNNs and the ensemble prediction accuracy per simulation end time.

Table B8 displays the neural network predictions for mutations from hydrophobic, non aromatic to hydrophobic,non aromatic amino acids. The WT is inserted as reference, to separate high from low binding affinity. H represents higher binding affinity and L represents lower binding affinity. The table lists the average accuracy of 10 CNNs and the ensemble prediction accuracy per simulation end time.

## Appendix C: Data availability

The distance matrices used to train the neural network are available at the following url: https://ibm.box.com/v/DistanceMatricesMD Supplementary videos are at the following url: https://ibm.box.com/v/SupplementaryVideos

## Appendix D: Code availability

The code used for data preprocessing and neural network trainings/predictions used in the study are available on github.com (https://github.com/CCCofficial/ML_spike_protein.git).

## Acknowledgements

We thank Binquan Luan for fruitful scientific discussion. We thank Alex Lenail for developing the online drawing tool called NN-SVG (https://alexlenail.me/NN-SVG/AlexNet.html). We used it to plot the architecture of the neural network we used in and L represents lower binding affinity.this study. We acknowledge support from the IBM Research AI Hardware Center, and the Center for Computational Innovation at Rensselaer Polytechnic Institute for computational resources on the AiMOS Supercomputer. This material is based upon work supported by the National Science Foundation under Grant No. DBI-1548297.

---

**Fig. A1.**
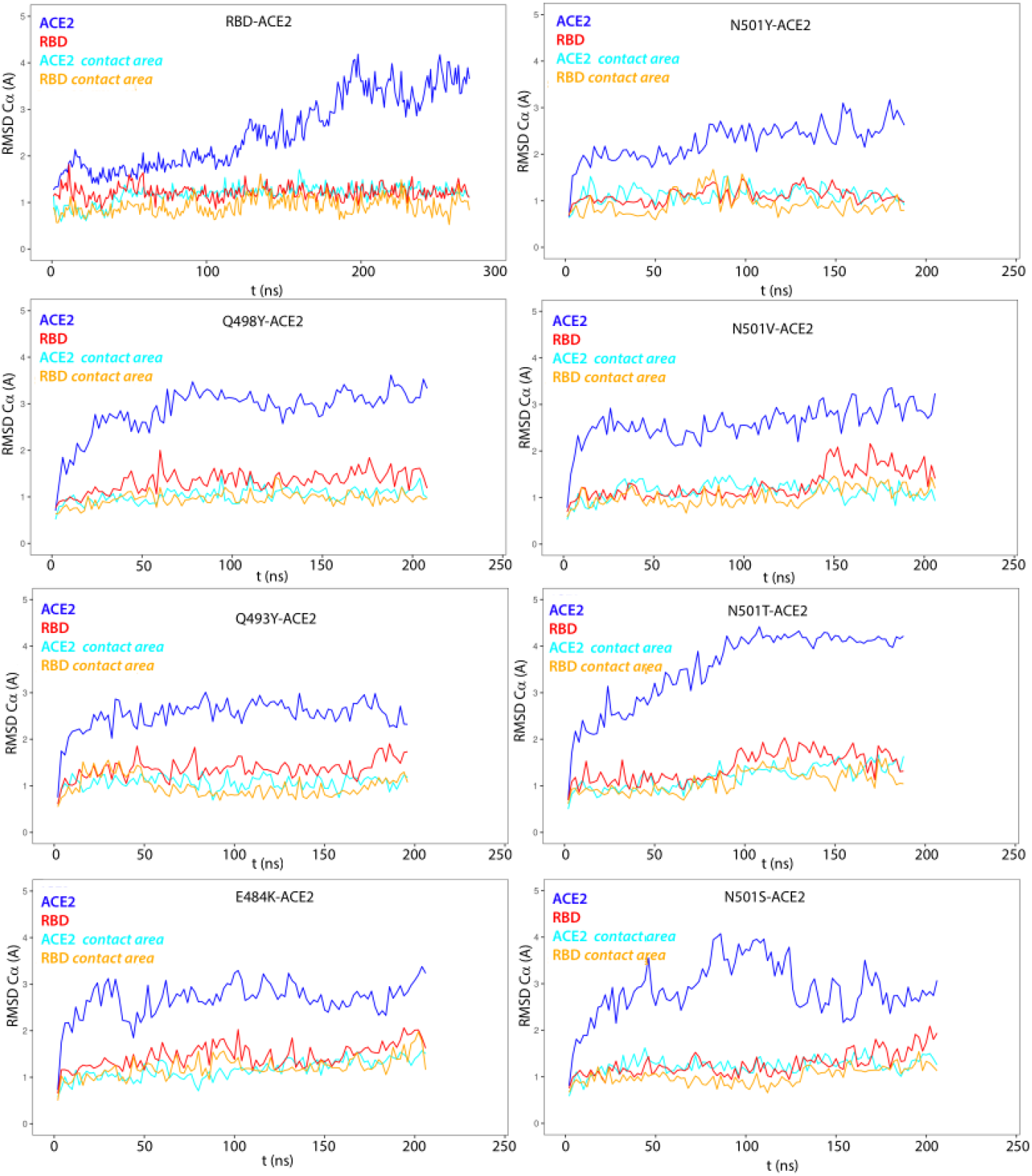
RMSD calculated for the C_*α*_ carbon atoms of the RBD and ACE2 and for the contact area for the simulations reported in the panels. For clarity, data are represented every 2 *ns*.

**Table B1.** Neural network predictions for Q498Y: This is the table listing the predictions from 10 neural networks for mutation Q498Y, which has higher affinity than the unmutated SARS-CoV-2 RBD. NN1 to NN10 represents the 10 neural networks we trained. FP is short for final prediction and PPC represents the percentage of CNNs predicting correctly. H represents higher binding affinity

| NNs | Simulation end time |  |  |  |  |  |  |  |
| --- | --- | --- | --- | --- | --- | --- | --- | --- |
|  | 20ns | 40ns | 60ns | 80ns | 100ns | 120ns | 140ns | 160ns |
| NN1 | H(51%) | H(64%) | H(69%) | H(88%) | H(63%) | H(94%) | H(84%) | H(69%) |
| NN2 | H(54%) | H(63%) | H(80%) | H(86%) | H(94%) | H(62%) | H(63%) | L(58%) |
| NN3 | L(64%) | H(75%) | H(99%) | H(87%) | H(70%) | H(69%) | H(78%) | H(78%) |
| NN4 | L(61%) | H(59%) | H(86%) | H(73%) | H(82%) | H(56%) | H(61%) | H(98%) |
| NN5 | L(64%) | H(74%) | H(73%) | L(61%) | H(84%) | H(64%) | H(89%) | H(85%) |
| NN6 | H(53%) | H(77%) | H(70%) | H(94%) | H(84%) | H(58%) | H(63%) | H(84%) |
| NN7 | H(51%) | L(62%) | H(54%) | H(84%) | H(97%) | H(94%) | H(60%) | H(77%) |
| NN8 | L(65%) | H(85%) | H(84%) | H(60%) | H(83%) | H(86%) | H(84%) | H(61%) |
| NN9 | H(51%) | H(80%) | H(70%) | H(60%) | H(81%) | H(57%) | L(60%) | H(65%) |
| NN10 | H(69%) | L(74%) | H(89%) | H(93%) | H(64%) | H(99%) | H(80%) | H(69%) |
| FP | H | H | H | H | H | H | H | H |
| PPC | 60% | 80% | 100% | 90% | 100% | 100% | 90% | 90% |

**Fig. A2.**
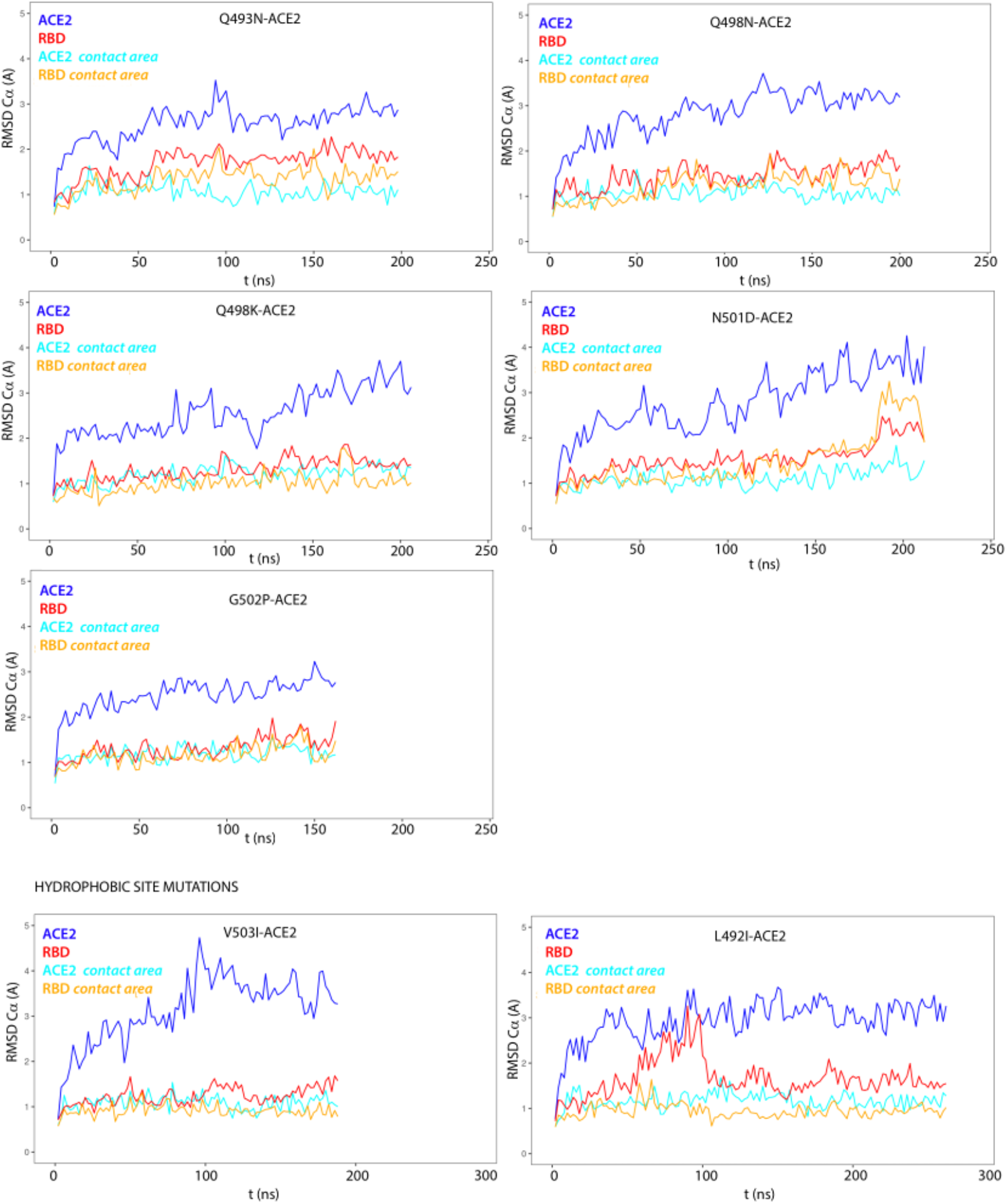
RMSD calculated for the C_*α*_ carbon atoms of the RBD and ACE2 and for the contact area for the simulations reported in the panels. For clarity, data are represented every 2*ns*.

**Fig. A3.**
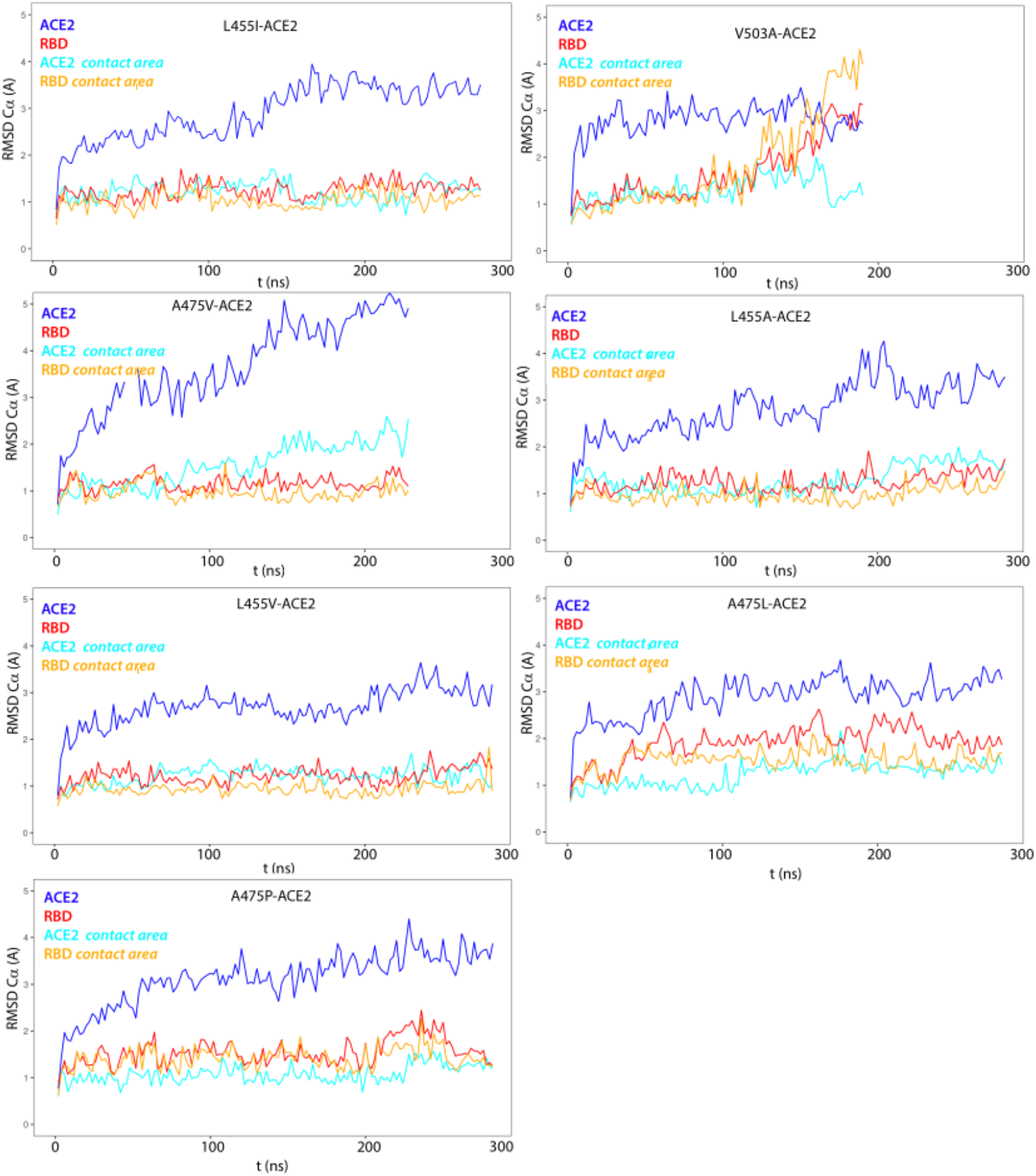
RMSD calculated for the C_*α*_ carbon atoms of the RBD and ACE2 and for the contact area for the simulations reported in the panels. For clarity, data are represented every 2*ns*.

**Table B2.** Neural network predictions for Q493Y: This is the table listing the predictions from 10 neural networks for mutation Q493Y, which has higher affinity than the unmutated SARS-CoV-2 RBD. *The number of H is the same as the number of L. The final prediction is based on comparing the averaged *P*_1_ (=0.854) for H and averaged *P*_2_ (= 0.628) for L and choosing the larger one. NN1 to NN10 represents the 10 neural networks we trained. FP is short for final prediction and PPC represents the percentage of CNNs predicting correctly. H represents higher binding affinity and L represents lower binding affinity.

| NNs | Simulation end time |  |  |  |  |  |  |  |
| --- | --- | --- | --- | --- | --- | --- | --- | --- |
|  | 20ns | 40ns | 60ns | 80ns | 100ns | 120ns | 140ns | 160ns |
| NN1 | H(100%) | L(59%) | H(99%) | H(67%) | H(71%) | H(96%) | H(86%) | H(97%) |
| NN2 | H(78%) | L(64%) | H(98%) | H(99%) | H(100%) | H(82%) | H(99%) | H(83%) |
| NN3 | H(74%) | H(63%) | H(84%) | H(69%) | H(70%) | H(98%) | H(99%) | H(100%) |
| NN4 | H(87%) | L(67%) | H(97%) | H(97%) | H(99%) | H(75%) | H(86%) | H(100%) |
| NN5 | H(60%) | L(67%) | H(51%) | H(78%) | H(95%) | H(87%) | H(99%) | H(97%) |
| NN6 | H(85%) | H(99%) | H(95%) | H(96%) | H(68%) | H(99%) | H(99%) | H(98%) |
| NN7 | H(99%) | L(57%) | H(70%) | H(97%) | H(99%) | H(91%) | H(99%) | H(99%) |
| NN8 | H(79%) | H(82%) | H(58%) | H(72%) | H(97%) | H(99%) | H(100%) | H(92%) |
| NN9 | H(90%) | H(85%) | L(53%) | H(75%) | H(95%) | H(56%) | H(80%) | H(94%) |
| NN10 | H(84%) | H(98%) | L(58%) | H(95%) | H(76%) | H(100%) | H(97%) | H(82%) |
| FP | H | H* | H | H | H | H | H | H |
| PPC | 100% | 50% | 80% | 100% | 100% | 100% | 100% | 100% |

**Table B3.**
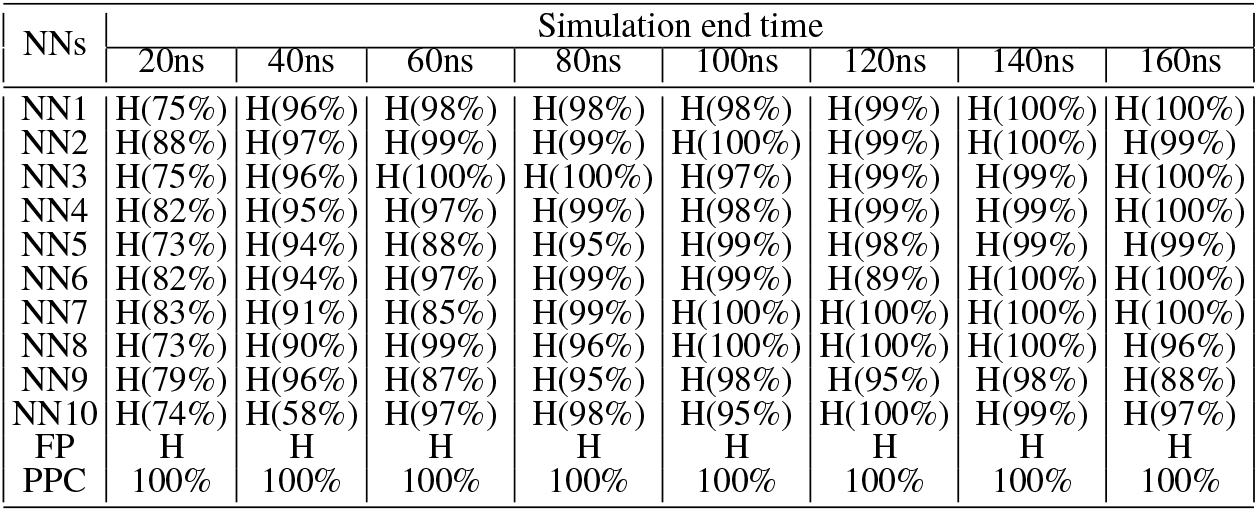
Neural network predictions for E484K: This is the table listing the predictions from 10 neural networks for mutation E484K, which has higher affinity than the unmutated SARS-CoV-2 RBD. NN1 to NN10 represents the 10 neural networks we trained. FP is short for final prediction and PPC represents the percentage of CNNs predicting correctly. H represents higher binding affinity and L represents lower binding affinity.

**Table B4.**
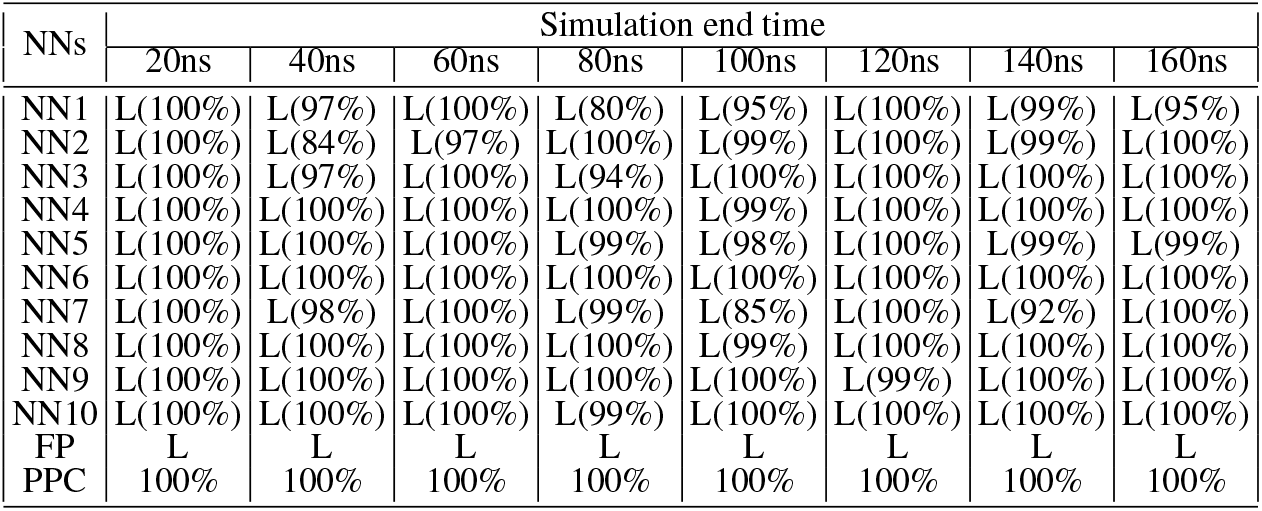
Neural network predictions for Q493N: This is the table listing the predictions from 10 neural networks for mutation Q493N, which has lower affinity than the unmutated SARS-CoV-2 RBD. NN1 to NN10 represents the 10 neural networks we trained. FP is short for final prediction and PPC represents the percentage of CNNs predicting correctly. H represents higher binding affinity and L represents lower binding affinity.

**Table B5.** Neural network predictions for Q498K: This is the table listing the predictions from 10 neural networks for mutation Q498K, which has lower affinity than the unmutated SARS-CoV-2 RBD. NN1 to NN10 represents the 10 neural networks we trained. FP is short for final prediction and PPC represents the percentage of CNNs predicting correctly. H represents higher binding affinity and L represents lower binding affinity.

| NNs | Simulation end time |  |  |  |  |  |  |  |
| --- | --- | --- | --- | --- | --- | --- | --- | --- |
|  | 20ns | 40ns | 60ns | 80ns | 100ns | 120ns | 140ns | 160ns |
| NN1 | H(68%) | L(85%) | L(72%) | L(77%) | H(60%) | L(76%) | H(54%) | L(75%) |
| NN2 | H(56%) | L(69%) | L(70%) | L(81%) | H(56%) | L(66%) | L(68%) | H(61%) |
| NN3 | H(78%) | L(78%) | L(56%) | L(65%) | L(87%) | L(84%) | L(77%) | L(66%) |
| NN4 | H(89%) | L(59%) | L(60%) | L(67%) | L(68%) | L(63%) | L(71%) | L(62%) |
| NN5 | L(66%) | L(64%) | L(57%) | H(88%) | L(77%) | L(76%) | L(70%) | L(53%) |
| NN6 | H(90%) | L(84%) | L(70%) | L(66%) | L(71%) | L(93%) | H(63%) | L(55%) |
| NN7 | H(90%) | L(82%) | L(77%) | L(73%) | H(71%) | L(56%) | L(56%) | L(57%) |
| NN8 | H(80%) | L(57%) | L(77%) | L(83%) | L(58%) | L(72%) | H(54%) | L(68%) |
| NN9 | H(78%) | H(64%) | L(74%) | L(83%) | L(57%) | H(69%) | L(80%) | L(64%) |
| NN10 | H(80%) | L(58%) | L(74%) | L(87%) | L(76%) | L(55%) | L(67%) | L(70%) |
| FP | H | L | L | L | L | L | L | L |
| PPC | 10% | 90% | 100% | 90% | 70% | 90% | 70% | 90% |

**Table B6.**
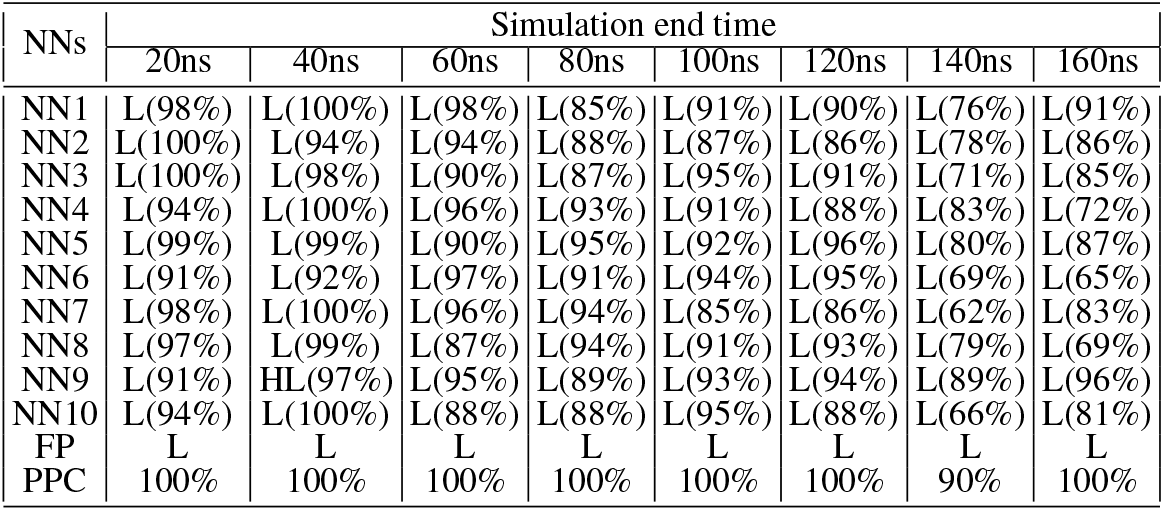
Neural network predictions for G502P: This is the table listing the predictions from 10 neural networks for mutation G502P, which has lower affinity than the unmutated SARS-CoV-2 RBD. NN1 to NN10 represents the 10 neural networks we trained. FP is short for final prediction and PPC represents the percentage of CNNs predicting correctly. H represents higher binding affinity and L represents lower binding affinity.

| NNs | Simulation end time |  |  |  |  |  |  |  |
| --- | --- | --- | --- | --- | --- | --- | --- | --- |
|  | 20ns | 40ns | 60ns | 80ns | 100ns | 120ns | 140ns | 160ns |
| NN1 | L(98%) | L(100%) | L(98%) | L(85%) | L(91%) | L(90%) | L(76%) | L(91%) |
| NN2 | L(100%) | L(94%) | L(94%) | L(88%) | L(87%) | L(86%) | L(78%) | L(86%) |
| NN3 | L(100%) | L(98%) | L(90%) | L(87%) | L(95%) | L(91%) | L(71%) | L(85%) |
| NN4 | L(94%) | L(100%) | L(96%) | L(93%) | L(91%) | L(88%) | L(83%) | L(72%) |
| NN5 | L(99%) | L(99%) | L(90%) | L(95%) | L(92%) | L(96%) | L(80%) | L(87%) |
| NN6 | L(91%) | L(92%) | L(97%) | L(91%) | L(94%) | L(95%) | L(69%) | L(65%) |
| NN7 | L(98%) | L(100%) | L(96%) | L(94%) | L(85%) | L(86%) | L(62%) | L(83%) |
| NN8 | L(97%) | L(99%) | L(87%) | L(94%) | L(91%) | L(93%) | L(79%) | L(69%) |
| NN9 | L(91%) | HL(97%) | L(95%) | L(89%) | L(93%) | L(94%) | L(89%) | L(96%) |
| NN10 | L(94%) | L(100%) | L(88%) | L(88%) | L(95%) | L(88%) | L(66%) | L(81%) |
| FP | L | L | L | L | L | L | L | L |
| PPC | 100% | 100% | 100% | 100% | 100% | 100% | 90% | 100% |

**Table B7.** Table listing the accuracy of the 10 independently trained neural network per simulation duration, i.e., we have trained 80 neural networks in total. This table also lists the average neural network accuracy and accuracy of ensemble predictions.

| Accuracy | Simulation end time |  |  |  |  |  |  |  |
| --- | --- | --- | --- | --- | --- | --- | --- | --- |
|  | 20ns | 40ns | 60ns | 80ns | 100ns | 120ns | 140ns | 160ns |
| NN1 | 5/6 | 5/6 | 6/6 | 6/6 | 5/6 | 6/6 | 5/6 | 6/6 |
| NN2 | 5/6 | 5/6 | 6/6 | 6/6 | 5/6 | 6/6 | 6/6 | 4/6 |
| NN3 | 4/6 | 6/6 | 6/6 | 6/6 | 6/6 | 6/6 | 6/6 | 6/6 |
| NN4 | 4/6 | 5/6 | 6/6 | 6/6 | 6/6 | 6/6 | 6/6 | 6/6 |
| NN5 | 5/6 | 5/6 | 6/6 | 4/6 | 6/6 | 6/6 | 6/6 | 6/6 |
| NN6 | 5/6 | 6/6 | 6/6 | 6/6 | 6/6 | 6/6 | 5/6 | 6/6 |
| NN7 | 5/6 | 4/6 | 6/6 | 6/6 | 5/6 | 6/6 | 5/6 | 6/6 |
| NN8 | 4/6 | 6/6 | 6/6 | 6/6 | 6/6 | 6/6 | 5/6 | 6/6 |
| NN9 | 5/6 | 5/6 | 5/6 | 6/6 | 6/6 | 5/6 | 5/6 | 6/6 |
| NN10 | 5/6 | 5/6 | 5/6 | 6/6 | 6/6 | 6/6 | 6/6 | 6/6 |
| Averaged NN prediction | 0.7833 | 0.8667 | 0.9667 | 0.9667 | 0.95 | 0.9833 | 0.9167 | 0.9667 |
| Ensemble Prediction | 5/6 | 1 | 1 | 1 | 1 | 1 | 1 | 1 |

**Table B8.** Neural network predictions for mutations from hydrophobic to hydrophobic amino acids. The WT is inserted as reference, to separate high from low binding affinity. H represents higher binding affinity and L represents lower binding affinity.

| Mutation | Neural network |  |  |  |  |  |  |  |  |  |  |
| --- | --- | --- | --- | --- | --- | --- | --- | --- | --- | --- | --- |
|  | NN1 | NN2 | NN3 | NN4 | NN5 | NN6 | NN7 | NN8 | NN9 | NN10 | Ensemble |
| V503I | H(59%) | L(57%) | H(62%) | H(84%) | H(58%) | H(52%) | H54(%) | L(60%) | L(52%) | L(64%) | H |
| L492I | L(54%) | L(60%) | H(50.4%) | L(63%) | L(62%) | L(60%) | L(62%) | L(63%) | L(79%) | L(66%) | L |
| WT |  |  |  |  |  |  |  |  |  |  |  |
| L455I | L(52%) | L(81%) | L(62%) | L(55%) | L(81%) | H(60%) | L(55%) | L(68%) | L(56%) | L(70%) | L |
| V503A | L(68%) | L(64%) | L(68%) | L(65%) | L(68%) | L(66%) | L(66%) | L(68%) | L(69%) | L(69%) | L |
| A475V | H(99.77%) | H(98%) | H(99.86%) | H(99.76%) | H(99%) | H(100%) | H(99.55%) | H(99%) | H(98%) | H(93%) | H |
| L455A | H(96%) | H(99%) | H(98%) | H(97%) | H(98%) | H(97%) | H(99%) | H(96%) | H(51%) | H(98%) | H |
| L455V | H(100%) | H(100%) | H(100%) | H(100%) | H(100%) | H(100%) | H(100%) | H(100%) | H(100%) | H(100%) | H |
| A475L | H(99%) | H(95%) | H(99.69%) | H(99%) | H(95%) | H(99%) | H(99%) | H(95%) | H(84%) | H(95%) | H |
| A475P | H(93%) | H(97%) | H(99.96%) | H(100%) | H(99%) | H(99%) | H(99%) | H(98%) | H(92%) | H(91%) | H |

